# REPLAY: A reproducible and user-friendly application for DNA replication timing analysis from Repli-seq data

**DOI:** 10.64898/2026.04.16.719037

**Authors:** Quinn Dickinson, Chuanhe Yu, Juan Carlos Rivera-Mulia

**Affiliations:** Department of Biochemistry, Molecular Biology and Biophysics, University of Minnesota Medical School, Minneapolis, Minnesota; Hormel Institute, University of Minnesota, Austin, Minnesota; Stem Cell Institute, University of Minnesota Medical School, Minneapolis, Minnesota; Masonic Cancer Center, University of Minnesota Medical School, Minneapolis, Minnesota; Masonic Institute on the Biology of Aging and Metabolism, University of Minnesota Medical School, Minneapolis, Minnesota

**Keywords:** DNA replication timing, Repli-seq, genomics, epigenome, automated workflow, Apptainer, reproducibility

## Abstract

**Background:** DNA replication timing (RT) is a fundamental feature of genome organization that is regulated in a cell-type-specific manner and frequently altered in disease. Repli-seq is the standard approach for genome-wide RT profiling; however, its analysis typically requires multiple independent tools and custom scripts, limiting reproducibility, portability, and accessibility, particularly for users without computational expertise. In addition, existing workflows often lack standardization and require substantial user intervention.

**Results:** We developed REPLAY, a fully automated, reproducible, and user-friendly application for replication timing analysis. REPLAY is distributed as a standalone executable that enables end-to-end processing from compressed FASTQ files to genome-wide RT profiles without requiring software installation or programming experience. Through an intuitive graphical interface, users can configure analysis parameters, including input and output directories, reference genome, normalization strategy (quantile, median, or interquartile range), and smoothing. The application integrates all processing steps—quality control, trimming, alignment, binning, RT log2 calculation, normalization, smoothing, and visualization— within a single automated workflow. Application of REPLAY to publicly available datasets demonstrate accurate reconstruction of RT profiles and high reproducibility across samples.

**Conclusions:** REPLAY offers a portable, reproducible, and accessible solution for the analysis of RT data. By eliminating the need for command-line tools and complex installations, it lowers the entry barrier enabling standardized analysis across diverse research settings.

## BACKGROUND

DNA replication in eukaryotic cells is a highly coordinated process that follows a defined temporal program known as replication timing (RT), where distinct genomic regions replicate at specific stages of S-phase [1–4]. This program is fundamentally linked to the functional organization of the nucleus, strongly correlated with 3D genome architecture, chromatin epigenetic states, and transcriptional activity [3, 5–8]. RT is highly cell-type specific, with approximately half of the genome undergoing dynamic reorganization during development to coordinate with cell fate specification [9–11]. Furthermore, aberrations in the RT program are linked to genome instability [12–16] and are a hallmark of disease, such as cancer [17–21], highlighting the importance of RT for genomic function.

The current gold standard for genome-wide RT profiling is Repli-seq [21–24]. In the standard “Early/Late” (E/L) Repli-seq, cells are pulse-labeled with a nucleotide analog (typically BrdU), sorted into early and late S-phase fractions via fluorescence-activated cell sorting (FACS), and processed for high-throughput sequencing [21, 22, 25, 26]. The resulting data is analyzed to generate a log2 ratio of early-to-late enrichment, providing a comprehensive map of replication domains across the genome [22, 27]. Despite the widespread utility of Repli-seq, bioinformatic analysis remains a significant bottleneck. Existing workflows are often fragmented, requiring researchers to manually combine independent tools for quality control, adapter trimming, alignment, filtering, and normalization [28–30]. This process demands substantial expertise in Unix and R environments, creating a steep entry barrier for non-computational researchers. Moreover, the lack of standardized software environments often leads to limited portability and reproducibility across different computational platforms, as variations in tool versions and normalization strategies can introduce inconsistencies between studies.

To address these limitations, we developed REPLAY, a fully automated, reproducible, and portable application designed to streamline Repli-seq data analysis. REPLAY integrates the entire processing workflow—from raw FASTQ files to normalized RT profiles—in an executable application that exploits the Snakemake [31] workflow management system. To ensure absolute reproducibility across diverse computational environments, including local workstations and High-Performance Computing (HPC) clusters, the application is distributed via an integrated pyinstaller executable and an Apptainer (Singularity) container [32]. Furthermore, REPLAY features a user-friendly graphical user interface, allowing researchers to execute complex bioinformatic tasks without programming expertise. By providing a unified and accessible application, REPLAY facilitates the standardization of RT analysis and its broader adoption across the research community.

## IMPLEMENTATION

### Software architecture and application design

REPLAY is distributed as a standalone executable application, allowing users to perform complete RT analysis without installing dependencies or interacting with command-line tools. The application integrates all workflow components within a unified environment, simplifying deployment across different operating systems.

We implemented REPLAY using Snakemake, a Python-based workflow management system that ensures task execution order, parallelization, and re-entrancy [31]. The complete execution workflow is illustrated in **Supplementary Figure 1**. To overcome dependency conflicts and ensure reproducibility across distinct platforms, the entire Snakemake workflow is encapsulated within an Apptainer (formerly Singularity) container [32, 33]. This containerization approach isolates the software environment, ensuring that all necessary dependencies are included and that the workflow operates consistently irrespective of the underlying operating system. This methodology is particularly advantageous for execution on High-Performance Computing (HPC) clusters, as well as in local workstation computers. REPLAY implements a complete workflow that fully automates the Repli-seq data processing, including sequencing read trimming and filtering, alignment to the reference genome, data binning, calculation of RT values (Log2[E/L]), normalization and smoothing (**Figure 1A**). Users can execute the application without requiring administrative privileges or manually installing bioinformatics tools, thereby simplifying the deployment process. Moreover, our portable application includes a pyinstaller executable implementing a graphical user interface (GUI), developed using the PySide6 module (Qt for Python) for cross-platform GUI applications, that simplifies the workflow and facilitates the processing of Repli-seq data, specifically designed for users without programming knowledge (**Figure 1B**). This significantly lowers the barrier to adoption and broadens the application’s usability in diverse research settings.

**Figure 1.**
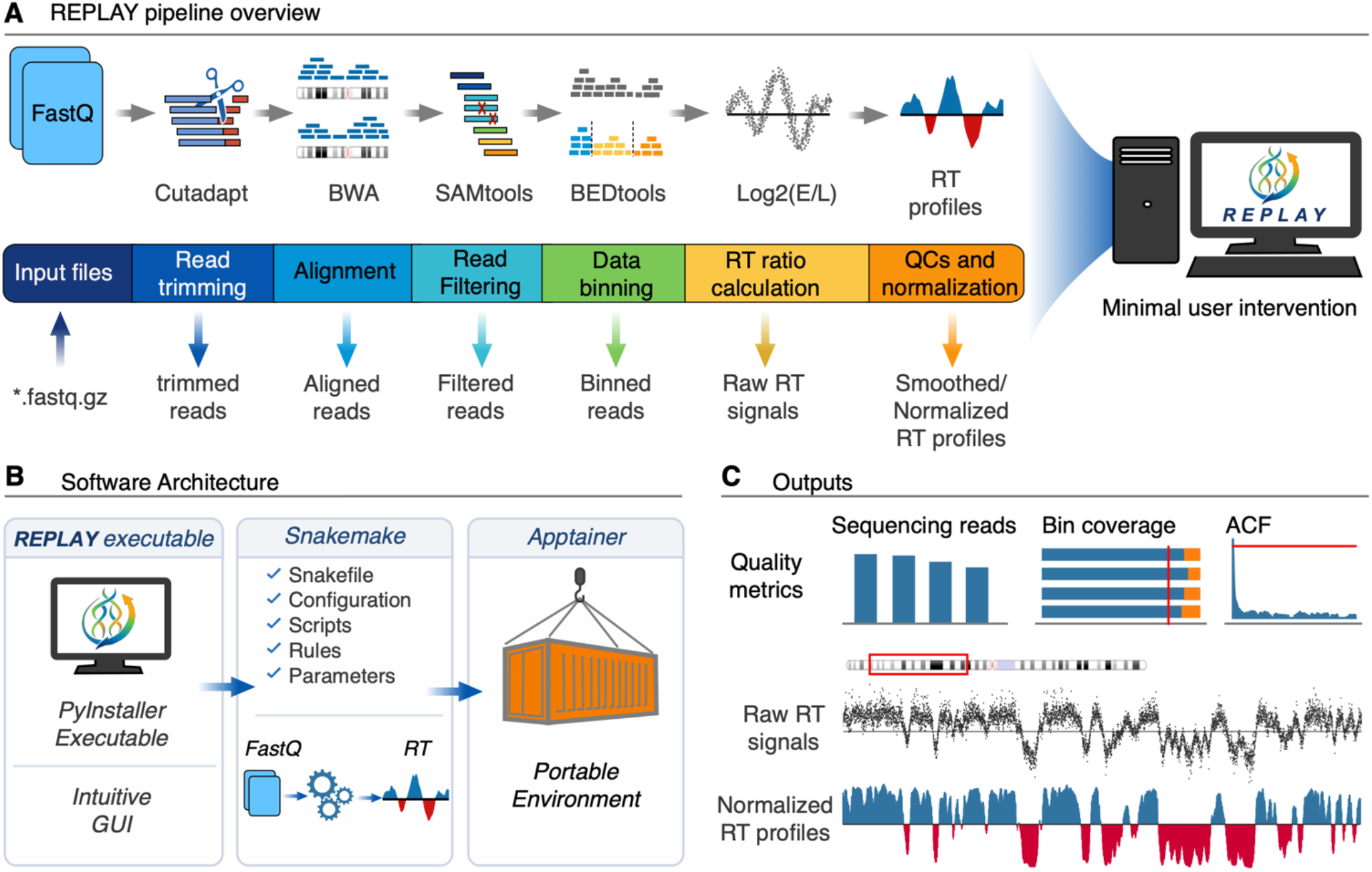
Overview of REPLAY application, from FASTQ to RT profiles. A) Schematic representation of the REPLAY software, illustrating the complete analysis workflow from compressed FASTQ files to RT profiles. Inputs consist of compressed FASTQ files from early and late S-phase fractions. Major steps include adapter trimming, alignment, read filtering and deduplication, genome binning, RT Log2 ratio calculation, normalization, smoothing, and visualization. B) Software architecture showing integration of Snakemake for efficient workflow management, friendly graphical user interface (GUI) and Apptainer containers for reproducible execution across computational environments. C) Summary of outputs include quality control metrics, raw, and normalized RT profiles.

The REPLAY automated workflow generates several key outputs. These outputs include the raw genome-wide RT profiles, which are expressed as RT Log2 ratios, as well as normalized and smoothed RT profiles. In addition to these profiles, the application produces quality and reproducibility metrics, essential for evaluating the reliability and accuracy of the generated data. The outputs, including the RT profiles and quality metrics, provide a clear and comprehensive overview of the analysis results (**Figure 1C**).

### Graphical user interface

REPLAY offers a user-friendly interface designed to simplify the entire process. The graphical user interface (GUI) was developed using PySide6 (Qt for Python) to enhance accessibility and user experience by configuring and executing the entire workflow without the need for complex command-line interactions. REPLAY-GUI allows users to easily select working and output directories, choose between single-end and pair-end sequencing types, select the reference genome and their preferred normalization and smoothing methods to suit their analysis needs (**Figure 2**). The platform accepts input files in the form of compressed FASTQ files (.*fastq*.*gz* extension). These files are automatically extracted and processed by our software, ensuring a seamless workflow. REPLAY interface is equipped with real-time progress monitoring and output visualization features, which facilitate workflow management and allow users to track the status of their tasks and view progress as they are processed (**Figure 2**). Additionally, our application automatically scans the files in the input directory, evaluates whether their names are valid for processing (including proper pairing of Early and Late S-phase libraries per sample), and renames the libraries if needed. Users can also process samples from distinct organisms by configuring their corresponding reference genomes, select the desired resolution (bin size), and filter the specific contig sequences to process (**Supplementary Figure 2**). Smoothing level is also customizable through the advanced settings (**Supplementary Figure 3**). Finally, the GUI also includes a “Results” section that compiles quality metrics including read mapping statistics, bin coverage, autocorrelation function, raw RT profile visualization, and raw and normalized data distributions (**Supplementary Figure 4**).

**Figure 2.**
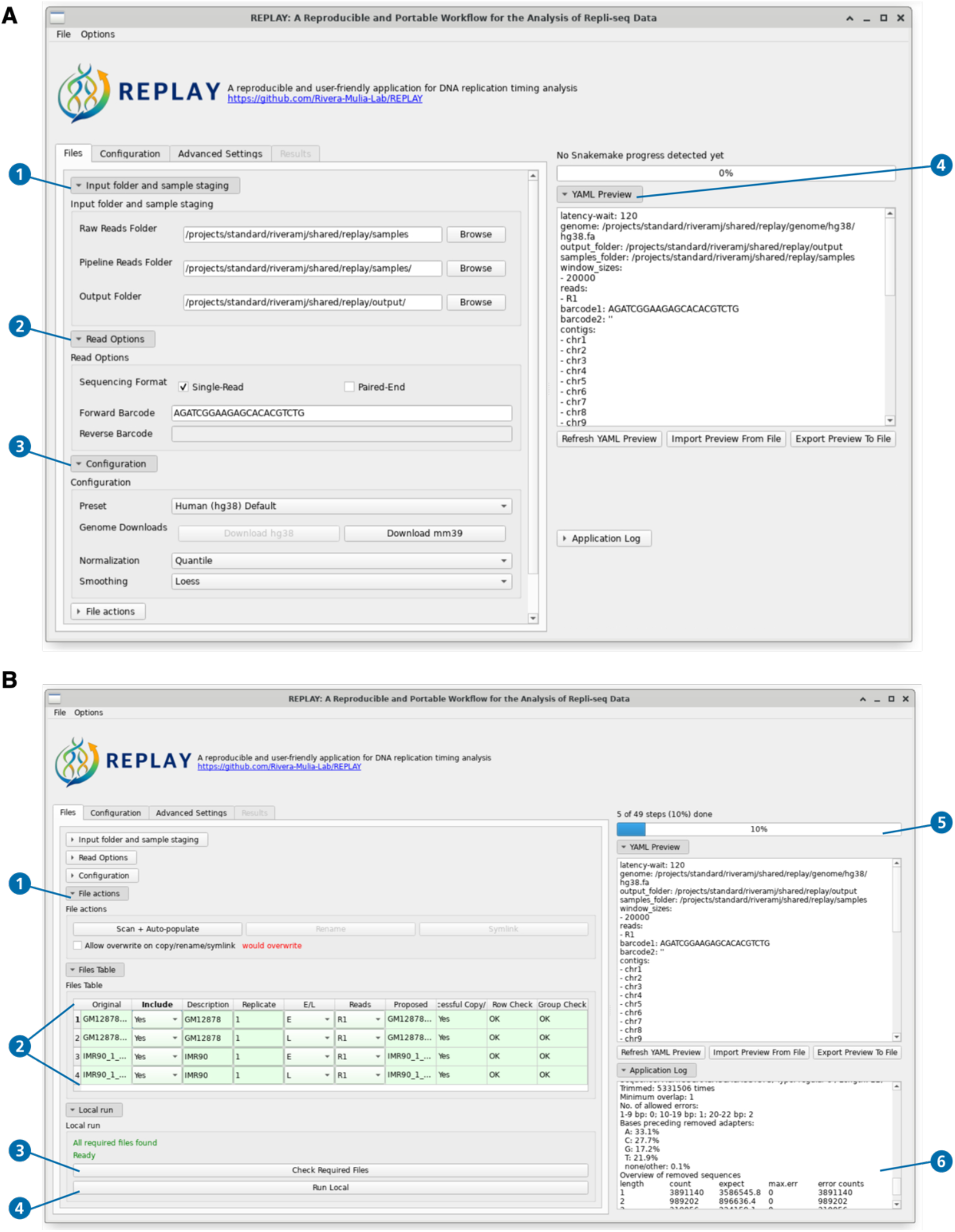
The standalone REPLAY application enables end-to-end analysis without command-line interaction. Screenshot of the REPLAY graphical user interface. A) The main window includes distinct collapsable menus: 1) selection for working and output directories; 2) selection of sequencing format (single or paired-ended sequencing); 3) configuration menu for selection of reference genome, RT data normalization strategy (quantile, interquartile, median, none), and smoothing (LOESS or Gaussian); 4) window containing the configuration parameters. B) A different view of the main window illustrating: 1) the files menu, which allow user to scan the input files to validate their names and rename them if necessary; 2) the list of detected files to be processed; 3) a check button to verify completeness of required files; 4) run button to initiate data processing; 5) progress bar for real-time monitoring of stepwise execution workflow; and 6) window of log files for tracking program execution.

### Data preprocessing and Alignment

The REPLAY application executes a series of pre-established steps to ensure accurate and efficient data processing:

1. Preprocessing: Adapter sequences are identified and removed. REPLAY removes adapters from raw *\**.*fastq*.*gz* files via cutadapt [34]. The cutadapt tool was selected because it supports distinct sequencing technologies, can remove sequences from both 5’ and 3’ ends, incorporates quality trimming, and supports pair-end reads. Moreover, it is commonly used for processing Repli-seq data [21, 22, 28, 29, 35]. REPLAY automatically populates the standard Illumina adapter sequences for single-end or pair-end sequencing data. However, users can input any custom adapter sequences.
2. Alignment: Following preprocessing, sequencing reads are mapped to the reference genome using the Burrows-Wheeler Alignment (BWA-MEM) tool [36]. This tool was chosen for its exceptional efficiency and accuracy in aligning sequencing reads from Repli-seq data [21, 22, 35].
3. Filtering: Post-alignment processing includes a filtering process to eliminate PCR duplicates and low-quality mappings, specifically those with a mapping quality score (MAPQ) below 30. This step is crucial for maintaining high-fidelity data. REPLAY utilizes SAMtools’ duplicate marking (markdup) feature [37], which was selected for its rapid performance and efficient memory usage.
4. Binning: The genome is partitioned into non-overlapping windows, with the size of these windows determined by user-defined resolutions, such as 5 kb, 10 kb, 20 kb, or 100 kb. REPLAY relies on BEDtools for genomic interval manipulation [38].
5. Quantification: In the final step, read counts are calculated per window and normalized as Reads Per Kilobase per Million mapped reads (RPKM). This normalization process ensures that the data is comparable across different samples and conditions, facilitating accurate quantitative analysis. The intermediary outputs are bedgraph files containing the reads per bin for each S-phase fraction, which are then processed in the subsequent steps for RT profile calculation.

### Calculation of RT log2 Ratios

RT profiles are determined by analyzing the relative abundance of DNA in early and late S-phase fractions of the cell cycle. To quantify this, REPLAY computes the RPKM per genomic bin in both early versus late S-phase fractions. Next, it calculates the ratio of RPKM values between these two phases. Finally, the software transforms these ratio values into a Log2 scale to facilitate easier comparison and interpretation of the data:

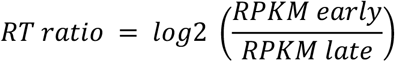

The resulting RT profiles are output as bedgraph files that contain the raw RT data across the genome, allowing for direct visualization and downstream analysis.

### Quality Control and Diagnostics

A key feature of REPLAY is the automated generation of comprehensive diagnostic metrics (**Figure 3**), which is critical for evaluating the integrity and reliability of the data being analyzed. These metrics evaluate data quality according to established field standards through several key analyses:

1. Read Mapping and Filtering Statistics: These statistics provide detailed information on the efficiency and accuracy of aligning sequence reads to the reference genome [21]. They help in understanding how well the reads are mapped and which reads are filtered out, ensuring that only high-quality data is retained for further analysis (**Figure 3A**).
2. Genome coverage Analysis: This analysis calculates the fraction of genomic windows that have non-zero coverage. For high-quality Repli-seq datasets, a threshold of ≥ 80% coverage is expected [21]. This ensures that a substantial portion of the genome is adequately sequenced, which is essential for accurate RT analyses (**Figure 3B**). Values lower than 80% indicate insufficient sequencing depth or poor S-phase fractions sorting.
3. Autocorrelation (ACF): The workflow computes the ACF value to assess the spatial consistency between adjacent replication domains [21, 26, 27]. Datasets that achieve an ACF value of ≥ 0.8 are flagged as high-quality (**Figure 3C**). This metric helps in determining the degree of correlation in the RT data, indicating how well the neighbor replication domains replicate consistently during the S-phase.
4. RT data visualization: REPLAY facilitates rapid visual inspection of the data. These visualizations include raw RT signals that are expected to clearly show enrichment towards early (positive values) and late (negative values) replication (**Figure 3D**). Raw RT signal distributions are also plotted to inspect potential bias in the data ((**Figure 3E**). High-quality Repli-seq datasets should show a bimodal distribution with peaks at early and late replication.

**Figure 3.**
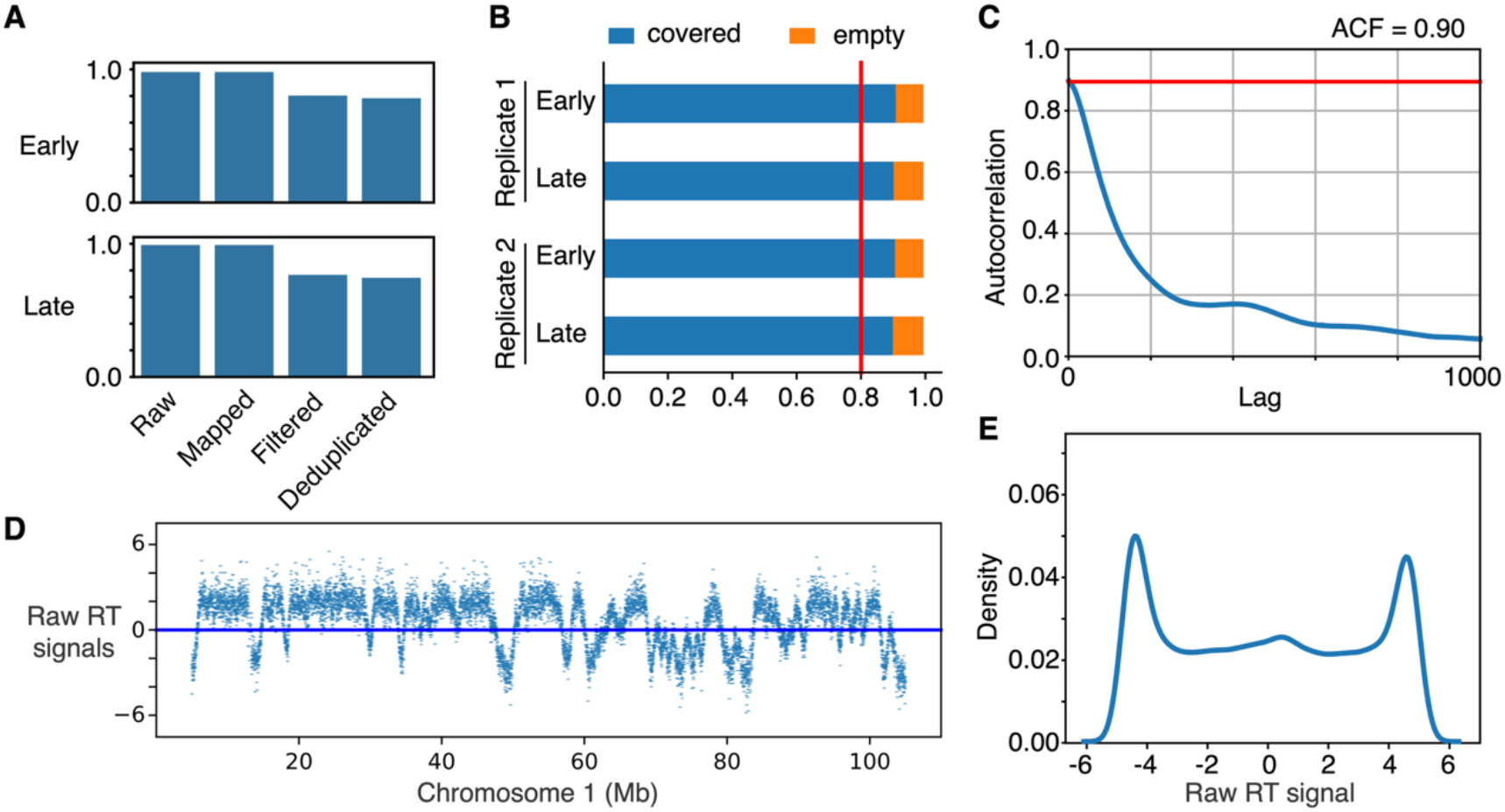
Quality control metrics generated by REPLAY. A) Summary of read processing across major application steps, including total reads, mapped reads, filtered reads, and deduplicated reads for early and late S-phase samples. B) Genome-wide coverage analysis showing fractions of bins containing reads and empty bins. A threshold of ≥0.8 is used to define high-quality sequenced libraries. C) Autocorrelation function (ACF) analysis of RT signal across genomic bins, illustrating spatial consistency in RT between adjacent replication domains. The decay of autocorrelation with increasing genomic distance reflects expected replication domain organization. D) Raw RT signal on chromosome 1. D) Distribution of raw RT signal. Repli-seq data from the IMR90 cell line from the 4D nucleome consortium [39] was used to generate the plots.

REPLAY compiles these quality metrics into a “Results” section for rapid user diagnostics (**Supplementary Figure 4**). These metrics collectively provide valuable insights into the quality of sequencing and the reliability of RT profiles, enabling researchers to make informed decisions regarding the data’s suitability for downstream analyses.

### Data normalization and smoothing

REPLAY provides distinct normalization strategies to address systematic biases and users can select the normalization method. These strategies include:

- None: No normalization is performed but data can still be smoothed to reduce noise in raw RT signals (see below).
- Quantile Normalization: This method adjusts the distribution of data points across various samples to achieve equivalent quantiles, thereby facilitating comparability. It is the recommended normalization technique commonly used for Repli-seq analyses [21, 22, 28, 29]. Users have the option to utilize the target dataset provided (generated by averaging multiple high-quality datasets) or select their own target dataset for normalization purposes.
- Median Normalization: This approach normalizes data by adjusting each sample’s median to a common value, thereby mitigating the impact of outliers.
- Interquartile Range (IQR)-based Normalization: This technique normalizes data by scaling it based on the interquartile range, which is the difference between the 75th and 25th percentiles. This approach effectively reduces the influence of extreme values on the data.

Additionally, REPLAY applies smoothing to further enhance data quality and reduce noise. One option is using Locally Estimated Scatterplot Smoothing (LOESS), a non-parametric approach that fits multiple regression models within localized subsets of the data [40]. This smoothing is performed separately for each chromosome, allowing the method to account for chromosome-specific variations. This LOESS smoothing interpolates the data through a low-degree polynomial fitting function to remove stochastic outliers. REPLAY uses a recommended setting of a span = 500 kilobases (kb). This window size is optimized for mammalian genomes to provide a flexible and robust way to smooth the signal while preserving a clear signal of early and late replication domains [21]. However, users can specify the desired window size to adjust the levels of smoothing (**Supplementary Figure 3**). Users may also select gaussian smoothing to apply a 1-D gaussian filter across the dataset or to apply no smoothing.

By applying these normalization and smoothing techniques, REPLAY provides RT profiles ready for further analysis and interpretation (**Figure 4**). Normalized and smoothed RT profiles are saved as bedGraph tracks, which can be readily visualized using standard tools or directly loaded in the Integrative Genomics Viewer (IGV) [41] or UCSC [42] browsers. This output format allows researchers to inspect and interpret the RT landscape across the genome, providing an accessible and intuitive way to verify data quality and identify biological patterns.

**Figure 4.**
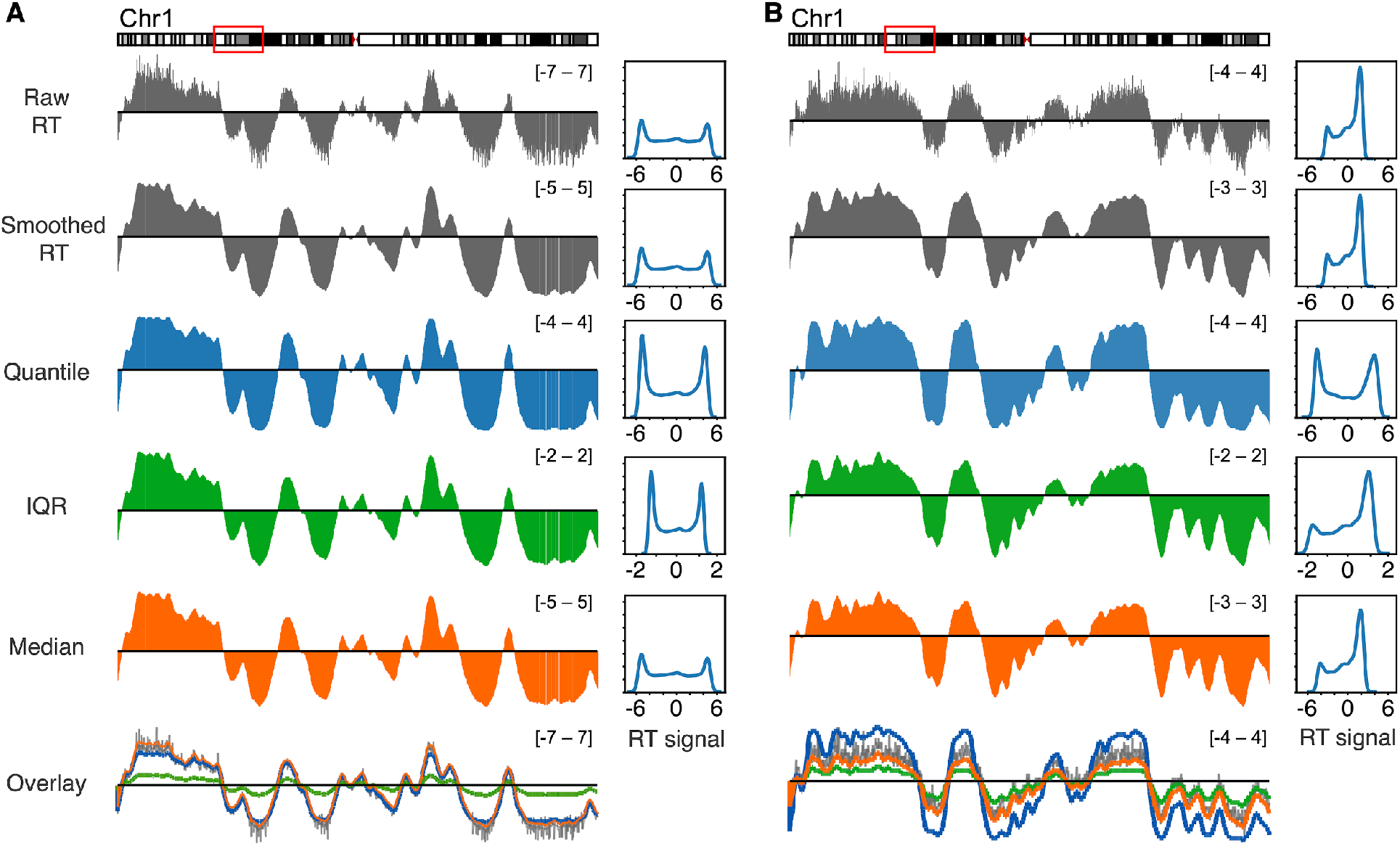
REPLAY normalization and smoothing of Repli-seq data. Raw RT profiles, smoothed, and normalized through distinct methods (quantile, interquartile, and median) are shown. Overlay plots and density distributions are shown to visualize the effects on the dynamic range using distinct normalization methods. Repli-seq data from the GM12878 (A) and IMR90 (B) cell lines obtained from the 4D nucleome consortium [39] were used.

### Reproducibility and software portability

Although distributed as an executable application, REPLAY internally leverages Snakemake [31] workflow management and a containerized environment to ensure reproducibility and consistent performance across platform systems. This ensures that the workflow can be adapted to different hardware and software configurations. All dependencies required for the workflow are encapsulated within an Apptainer container [32], providing a secure and portable way to package software and its dependencies, ensuring consistent behavior across different platforms and operating systems. This design choice allows REPLAY to be executed on both high-performance computing systems and local workstations, producing identical results from the same inputs provided (**Figure 5**).

**Figure 5.**
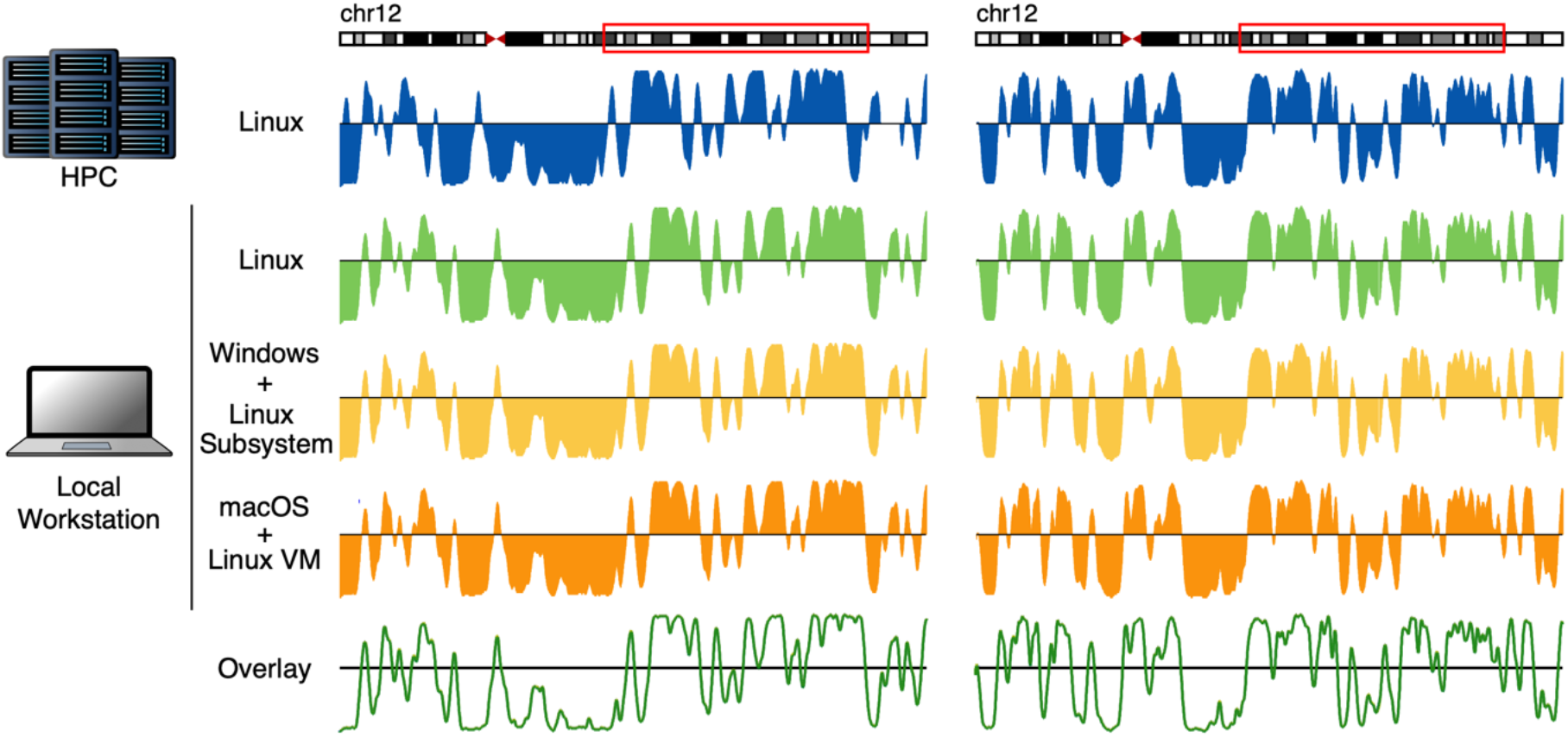
Reproducibility of REPLAY across computational environments and cell types. RT profiles generated by REPLAY across multiple computational environments and biological samples. RT profiles were computed from Repli-seq data for two human cell types, GM12878 (right) and IMR90 (left), and visualized across a representative chromosome. Repli-seq data was obtained from the 4D nucleome consortium [39]. Analyses were performed using REPLAY executed on (i) a high-performance computing environment at the Minnesota Supercomputing Institute, (ii) a Linux workstation, (iii) a Windows system using the Linux subsystem, and (iv) a macOS system with a virtualized Linux environment on Intel-based hardware. Across all platforms, REPLAY produced identical RT profiles, demonstrating consistent and reproducible behavior independent of computational environment.

To facilitate widespread adoption, REPLAY has been designed to operate seamlessly across all major operating systems (**Table 1**). The application is natively compiled for the x86-64 architecture, ensuring optimal compatibility with the server-grade hardware standard in genomic research. Windows users can launch REPLAY via the Windows Subsystem for Linux (WSL2), which provides a seamless, Linux-native execution experience through the graphical user interface. Similarly, REPLAY supports macOS on x86-64 (Intel) systems through virtualization (**Table 1**). Thus, REPLAY simplifies complex command-line-based bioinformatics tools by employing virtualization and containerization techniques.

**Table 1.** REPLAY cross-Platform Support Overview.

| Operating System | Implementation | User Experience |
| --- | --- | --- |
| Linux | Native Executable | Native: Double-click to launch; utilizes local Aptainer/Singularity. |
| Windows | Windows Subsystem for Linux (WSL2) | Runs within the WSL2 environment with full GUI support. |
| macOS (x86-64 architecture) | Linux Virtual Machine (VM) | Operates via VM (e.g., VirtualBox, Lima or Parallels) to bridge Linux-specific dependencies |

**Table 2.** Comparison of REPLAY with representative tools and workflows for Repli-seq data analysis. Features were assessed based on publicly available documentation and reported functionality.

|  | **REPLAY** | **ENCODE**<br>**Repli-seq**<br>**workflow** | **4D nucleome**<br>**Repli-seq**<br>**workflow** | **START-R** | **Repliseq-nf** |
| --- | --- | --- | --- | --- | --- |
| **End-to-end processing** | Complete<br>(FASTQ → RT<br>profiles) | No | No | No | Partial |
| **User interface** | Graphical user<br>interface (GUI) | Command line | Command line | Web-based<br>interface | Command<br>line |
| **Automation level** | Fully automated<br>(Snakemake) | Manual | Manual | Partial | Partial<br>(user-<br>configured) |
| **Workflow** | Fully automated<br>(Snakemake) | Multi-step<br>workflow (script-<br>based) | Multi-step<br>workflow (script-<br>based) | R Shiny App | Nextflow |
| **Programming**<br>**skills required** | No | Yes | Yes | No | Yes |
| **Containerization** | Apptainer | None | Docker | Docker | Docker/<br>Apptainer |
| **Input data** | Compressed<br>FASTQ | FASTQ | FASTQ | Preprocessed<br>BedGraph | FASTQ |
| **Output data** | Normalized and<br>smoothed RT,<br>QC metrics | Intermediate<br>signal tracks | Coverage /<br>RPKM signal | Normalized RT | Normalized<br>RT |
| **Normalization** | User-<br>configurable | Not included<br>(requires<br>downstream<br>processing) | Not included<br>(requires<br>downstream<br>processing) | User-<br>configurable | Quantile |
| **Smoothing** | User-<br>configurable | Not included<br>(requires<br>downstream<br>processing) | Not included<br>(requires<br>downstream<br>processing) | User-<br>configurable | User-<br>configurable |
| **Integrated QC**<br>**metrics** | Mapping,<br>coverage, ACF,<br>RT distributions | Standard<br>FASTQC | Not provided | Limited | Standard<br>FASTQC |
| **Primary use** | End-to-end<br>analysis | Data generation | Data generation | Visualization | Partial<br>pipeline |

## RESULTS

### Generation of RT profiles and validation

To assess the performance of REPLAY, we analyzed publicly available Repli-seq datasets generated across multiple human cell types. Specifically, we used datasets produced by the 4D Nucleome Project [39, 43], which provide high-quality measurements of early and late S-phase fractions. REPLAY successfully generated genome-wide RT profiles from raw sequencing data. To evaluate accuracy, we compared REPLAY-derived RT profiles to reference profiles computed using the 4D Nucleome processing pipeline [28]. This comparison demonstrated strong concordance across genomic regions (**Figure 6A–B**). In addition, analysis of independent biological replicates revealed high reproducibility, with RT profiles showing strong agreement across samples within the same cell type (**Figure 6C**). Distinct RT patterns observed between different cell types further confirm the ability of REPLAY to capture biologically meaningful variation in RT programs. Together, these results demonstrate that REPLAY accurately reconstructs genome-wide RT profiles and produces robust, reproducible outputs across datasets.

**Figure 6.**
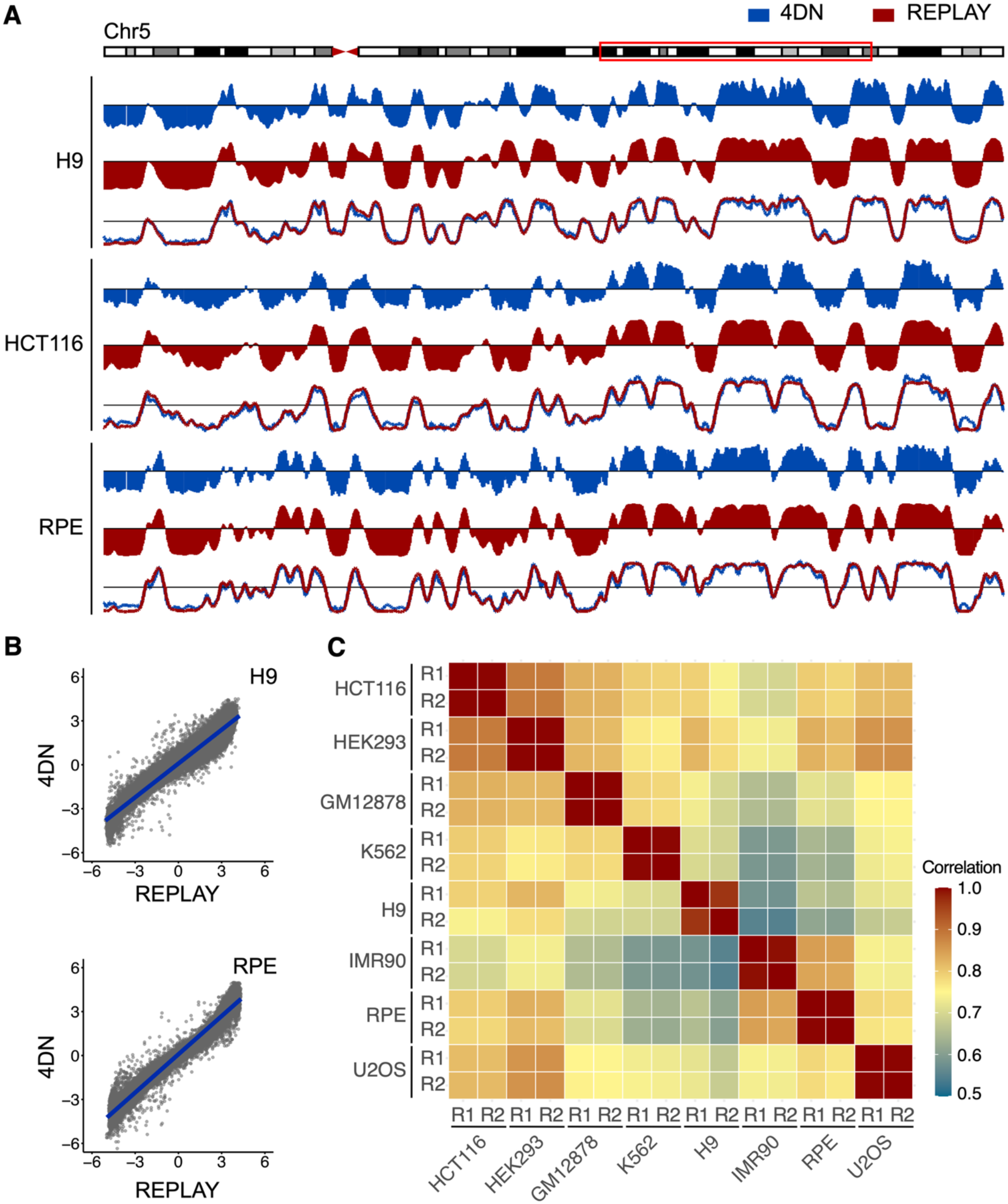
REPLAY generates accurate and reproducible RT profiles. A) RT profiles generated using the 4D Nucleome Project pipeline and REPLAY. Publicly available Repli-seq datasets from H9, HCT116, and RPE cell lines were analyzed. For each cell line, RT profiles are shown as individual tracks and as overlay plots, demonstrating concordance in domain structure and signal distribution. B) Genome-wide scatterplots comparing RT data generated using the 4D nucleome pipeline and REPLAY application. Data from H9 and RPE cell lines were used. A linear regression model is shown with slope approaching 1, indicating strong agreement between methods. C) Pearson correlation analysis of RT profiles processed with REPLAY across biological replicates. Repli-seq datasets from two replicates of HCT116, HEK293, GM12878, K562, H9, IMR90, RPE, and U2OS cell lines were analyzed at 20 kb resolution. High correlations (r ≥ 0.98) were observed across all replicate comparisons, demonstrating the reproducibility of REPLAY.

### Comparison with existing tools and workflow

Current approaches for Repli-seq data analysis rely on a combination of independent tools and partially automated workflows, often requiring substantial user intervention and computational expertise. Pipelines developed by large-scale efforts, including those from the ENCODE and the 4D Nucleome Project [28, 29], implement standardized processing steps such as read trimming, alignment, filtering, and signal generation. However, these workflows typically produce intermediate outputs (e.g., coverage or RPKM tracks) and do not include downstream steps such as normalization, smoothing, or final RT profile generation. As a result, additional processing and user intervention are required to obtain fully interpretable RT profiles. Other implementations from individual laboratories [44–50], are primarily command-line based and consist on multi-step workflows requiring user coordination across processing steps. Available tools greatly vary in scope and functionality. For example, START-R [30] provides an interactive web interface but is primarily designed for visualization, requires preprocessed genomic signal files, and does not support complete end-to-end processing of Repli-seq data. Workflow-based implementations such as Repli-seq-nf [50] improve automation but require manual configuration of parameters, command-line execution, and familiarity with workflow management systems. Because these tools vary in scope, level of automation, and required user intervention, direct benchmarking across methods is not straightforward. Thus, we compared their capabilities and level of integration (**Table 1**).

REPLAY distinguishes itself by integrating all steps of replication timing analysis within a single standalone application, enabling fully automated processing from raw FASTQ files to normalized RT profiles. Unlike traditional pipelines that require the manual setup of Conda environments, R libraries, and workflow managers, REPLAY is distributed as a standalone executable that includes all necessary dependencies. Through its graphical interface, users can configure parameters and execute analyses without command-line interaction or manual intervention. As summarized in **Table 1**, REPLAY is the only solution that combines end-to-end processing, full automation, and a graphical user interface within a unified framework.

## DISCUSSION

REPLAY addresses key limitations in current approaches to RT analysis by providing a unified, reproducible, and user-friendly application. Existing workflows typically rely on combinations of independent tools for quality control, alignment, and downstream processing, often requiring command-line execution and complex software environments [21, 22, 28, 29]. In contrast, REPLAY is distributed as a standalone executable that enables end-to-end analysis from raw FASTQ files to finalized RT profiles without requiring installation or programming expertise. By integrating all processing steps within a graphical interface, REPLAY reduces variability introduced by custom scripting and promotes reproducibility across laboratories.

A key feature of REPLAY is its level of integration and automation. While pipelines developed by large-scale efforts such as the ENCODE and the 4D Nucleome Project implement standardized preprocessing steps, they typically generate intermediate outputs (e.g., coverage or RPKM tracks) and do not include downstream normalization, smoothing, or final RT profile generation. As a result, additional processing and user intervention are required to obtain biologically interpretable RT profiles. Other tools address specific aspects of the workflow but differ in scope and level of automation, making direct benchmarking challenging. By contrast, REPLAY integrates all steps within a single framework, enabling fully automated and standardized analysis.

In addition to usability, REPLAY provides substantial improvements in computational efficiency and portability. End-to-end analyses are completed in less than one hour for a human sample, compared to substantially longer runtimes reported for multi-step workflows. These gains are achieved without compromising accuracy, as RT profiles generated by REPLAY show high concordance with independently derived profiles (Pearson’s r ≥ 0.98). Furthermore, containerization using Apptainer (Singularity) ensures a consistent computational environment, enabling reproducible results across platforms ranging from local workstations to high-performance computing (HPC) systems. The inclusion of a graphical user interface (GUI) implemented with PySide6 further expands accessibility. By allowing users to configure parameters such as bin size, normalization method, and smoothing without command-line interaction, REPLAY lowers the entry barrier for researchers without computational expertise and facilitates broader adoption of RT profiling.

The current version of REPLAY is optimized for standard early/late (E/L) Repli-seq data. Future developments will focus on extending this framework to support multi-fraction Repli-seq, enabling higher temporal resolution and improved detection of replication initiation and termination zones [35, 51]. In addition, extending REPLAY to single-cell Repli-seq data analysis will provide a standardized framework for investigating cell-to-cell variability in RT. This is particularly relevant given recent conflicting reports on the establishment of RT programs during early mammalian development [15, 52–54], highlighting the need for robust computational approaches that incorporate stringent quality control, minimize analytical biases, and identify technical artifacts that can confound the interpretation of single-cell RT data [55]. Finally, the modular and containerized design of REPLAY provides a foundation for integration with emerging multi-omic approaches, such as PARTAGE, which enables the joint profiling of copy number variation, replication timing, and gene expression from the same sample [35]. Together, these features position REPLAY as a comprehensive and accessible framework for standardized RT analysis across diverse experimental and computational settings.

## CONCLUSIONS

REPLAY provides an integrated and user-friendly solution for Repli-seq data analysis, bridging the gap between complex bioinformatics workflows and experimental research needs. By combining automated workflow management, containerized and portable software environments, and an intuitive graphical interface within a standalone application, REPLAY enables end-to-end generation of high-quality replication timing profiles with minimal user intervention. This framework facilitates reproducible and standardized analyses, supporting broader adoption of replication timing studies and improving consistency across datasets and laboratories.

## AVAILABILITY AND REQUIREMENTS

Project name: REPLAY

Project home page: https://github.com/Rivera-Mulia-Lab/REPLAY

Operating system(s): Linux, Windows (via WSL2), and macOS (via virtualization)

Programming language: Python, Bash.

Other requirements:

- Processor (CPU): x86-64 architecture. Minimum 2 cores. 6 cores or higher recommended. The Snakemake backend automatically parallelizes tasks based on available threads.
- Memory (RAM): 10 GB minimum. 32 GB or higher is recommended for processing large mammalian genomes (e.g., human or mouse) at high resolutions (< 10 kb bins).
- Storage:
  - Containers: ∼1 GB for the executable and Apptainer/Singularity image.
  - Genome + Index: ∼8.3 GB for hg38
  - Input Data: Variable (depends on FASTQ size).
  - Output Data: Assume at least ∼13x input for intermediate files. e.g. 1.21GB fastq.gz -> 14.8GB trimmed reads/bam/bedgraphs

License: GPL-3.0.

## AUTHOR CONTRIBUTIONS

Conceptualization and workflow design: Q.D. and J.C.R.M. Software implementation: Q.D. Writing original draft: J.C.R.M. and Q.D. Data analysis: Q.D. and J.C.R.M. Visualization: Q.D., and J.C.R.M. Study supervision: C.Y. and J.C.R.M.

## ACKNOWLEDGEMENTS

The authors acknowledge the Minnesota Supercomputing Institute (MSI) at the University of Minnesota for providing resources that contributed to the research results reported within this paper. During the preparation of this work the authors used GPT-5.3 in order to assist with the development, refinement and debugging of the included code. The authors reviewed and edited the code as needed and take full responsibility for the content of the publication.

