## Supplementary Information for "REPLAY: A reproducible and user-friendly application for DNA replication timing analysis from Repli-seq data"

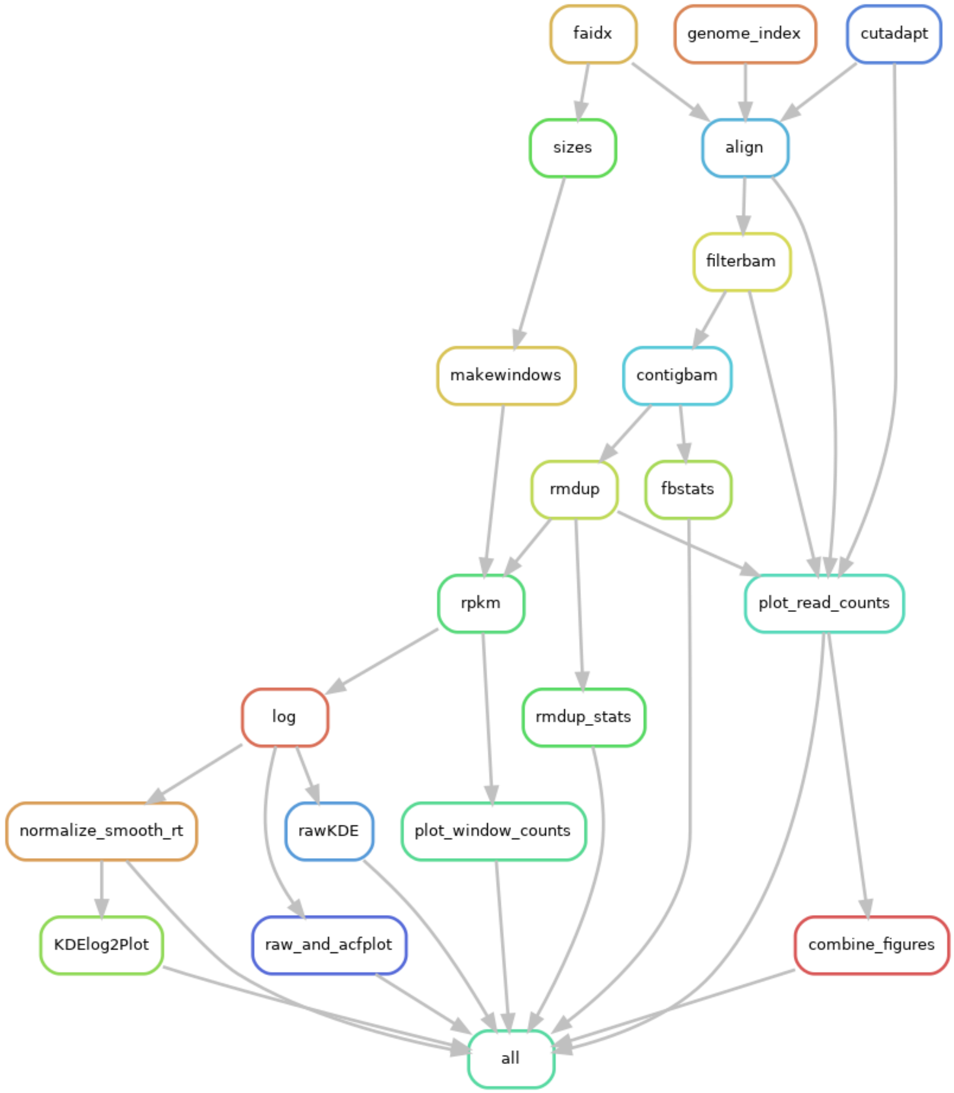


**Supplementary Figure 1. Snakemake workflow for replication timing analysis from raw sequencing reads to normalized profiles.** Directed acyclic graph (DAG) illustrating the full computational pipeline implemented in Snakemake, starting from compressed FASTQ files and ending with normalized and smoothed replication timing (RT) profiles and summary figures. Initial preprocessing steps include adapter trimming (cutadapt), genome indexing (genome_index, faidx), followed by read alignment (align). Aligned reads are subsequently processed through filtering (filterbam), BAM contig filtering (contigbam), duplicate removal (rmdup), and quality/statistics reporting (fbstats and rmdup_stats). Genome-wide binning is performed using generate windows (makewindows) in combination with chromosome size information (sizes), enabling downstream quantification of read coverage. Normalized signal extraction is performed via RPKM calculation (rpkm), which is further processed through log transformation (log) and smoothing/normalization steps (normalize_smooth_rt) to generate final replication timing profiles. Downstream analyses include window-based coverage visualization (plot_window_counts), read-level QC visualization (plot_read_counts), autocorrelation and raw signal assessment (rawKDE and raw_and_acfplot), and genome-wide smoothed profile visualization (KDElog2Plot). Final outputs are consolidated into the terminal target rule (all).

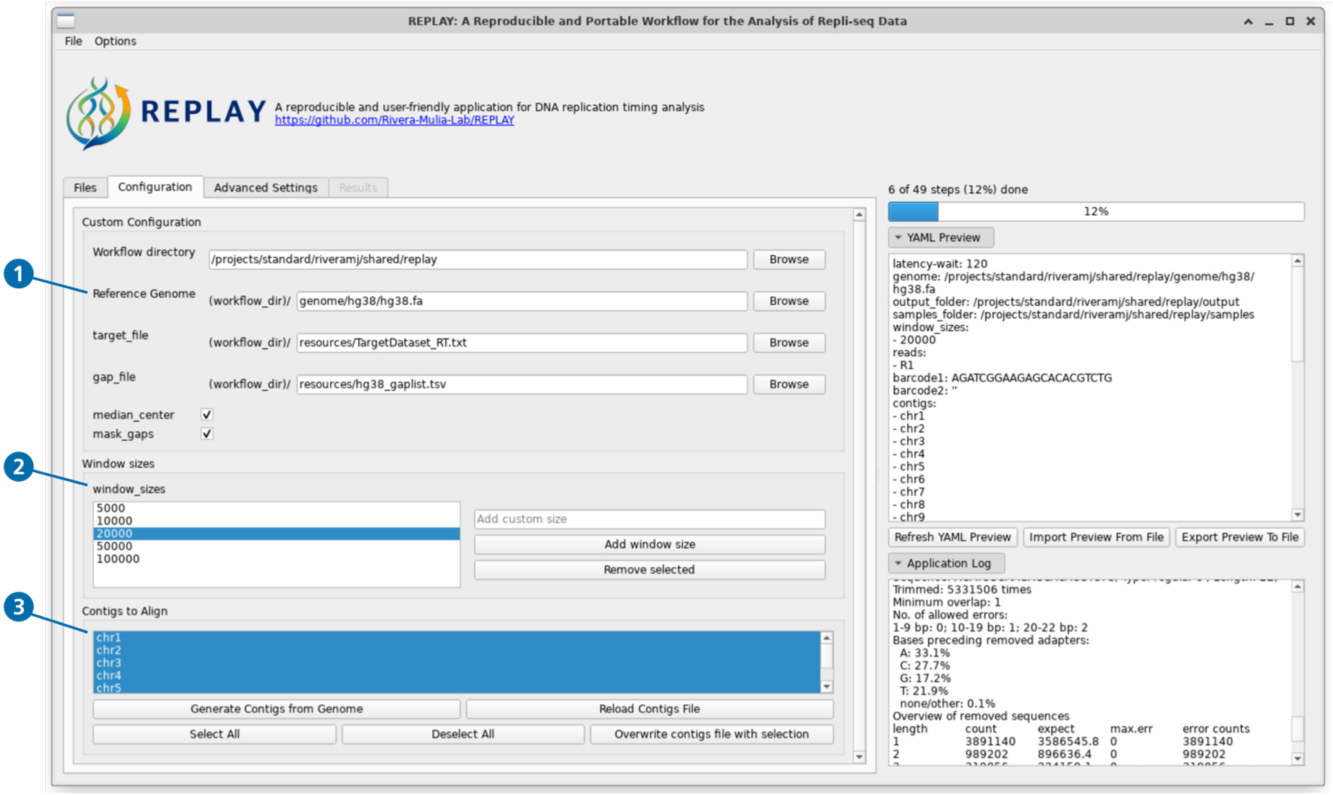


**Supplementary Figure 2.** Screenshot of the REPLAY configuration menu. 1) selection of reference genome to be used, which can be customized to any reference genome for distinct organisms; 2) selection options of RT profile resolutions (bin size); 3) specific chromosomal contig sequences to process, enabling the removal of alternative haplotypes, structural variants, or unplaced regions.

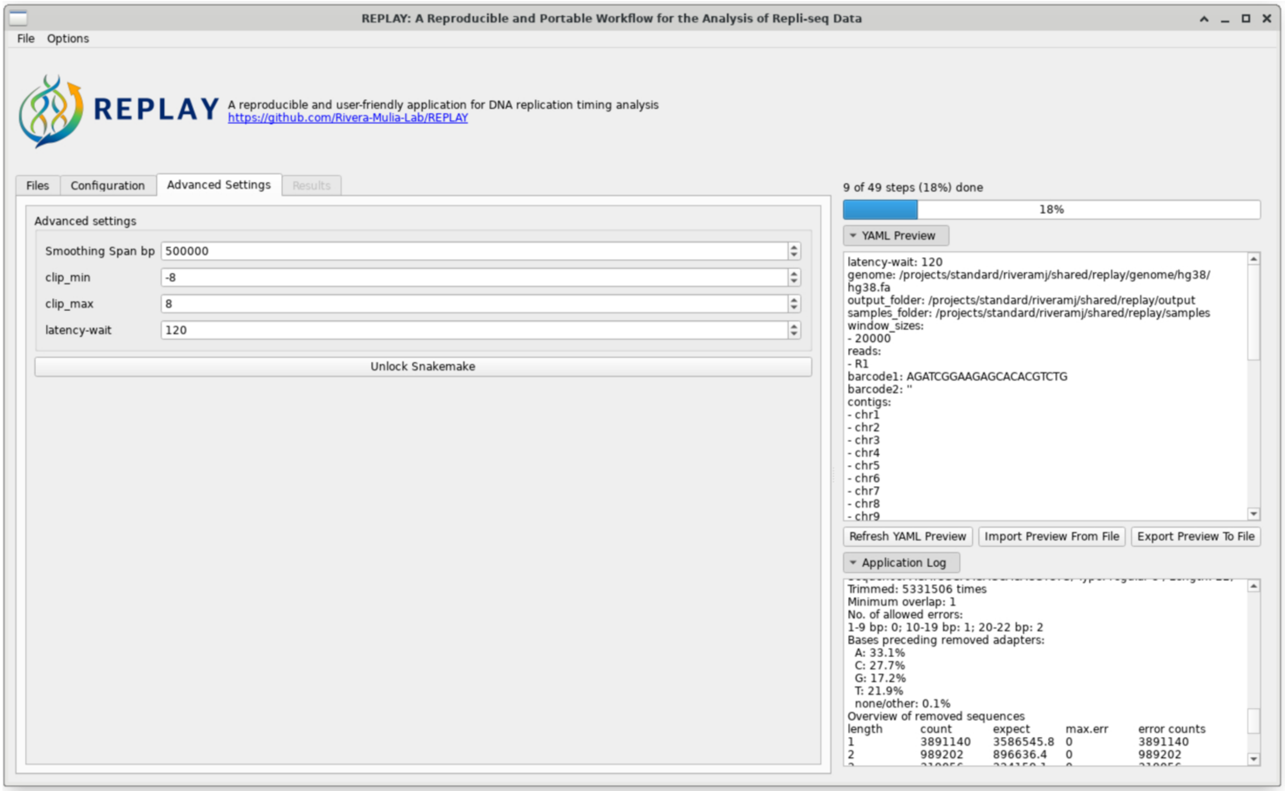


**Supplementary Figure 3**. Screenshot of the REPLAY Advanced settings menu. This menu enables the customization of the smoothing parameters and to unlock the Snakemake directory in the event of a crash.

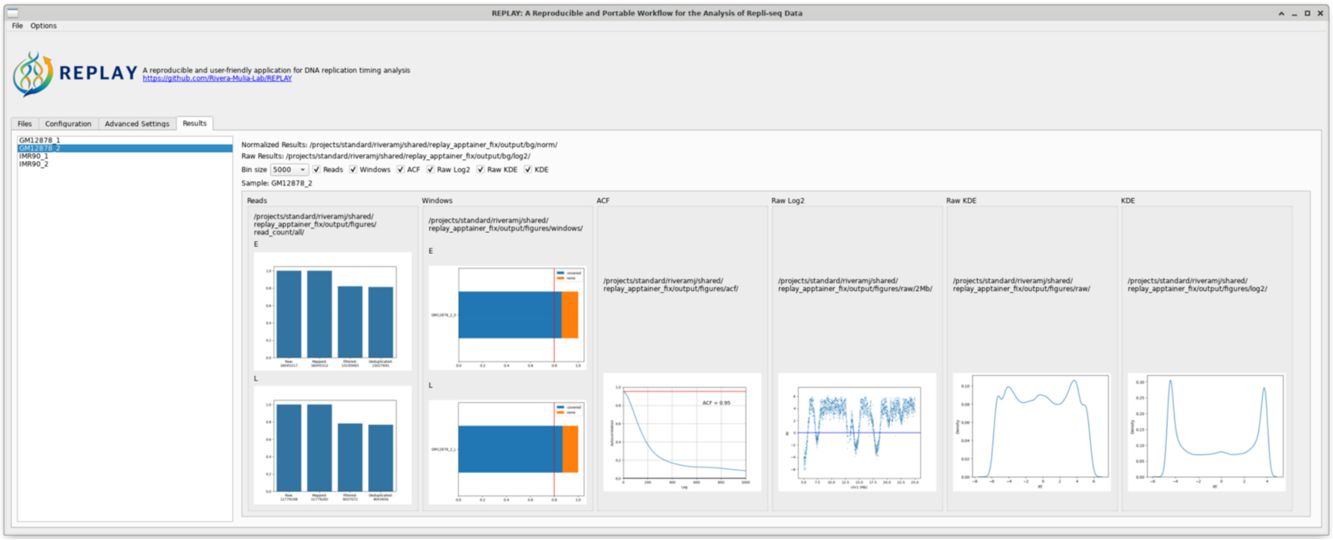


**Supplementary Figure 4**. Screenshot of the REPLAY Results window. This window compiles all results of the Repli-seq analysis process. Quality metrics include read mapping statistics, bin coverage, autocorrelation function, raw RT profile visualization, and raw and normalized data distributions
